# PIQUE: An ImageJ plugin for the quantification of toroidal nuclei in biological images

**DOI:** 10.1101/2022.11.07.515391

**Authors:** Carles Pons, Caroline Mauvezin

## Abstract

The toroidal nucleus is a novel chromosomal instability (CIN) biomarker which complements the micronucleus. Understanding the specific biological stresses leading to the formation of each CIN-associated phenotype requires the evaluation of large panels of biological images collected from different genetic backgrounds and environmental conditions. However, the quantification of toroidal nuclei is currently a manual process which is unviable on a large scale. Here, we present PIQUE (Processing of Images for the QUantification of toroidal nuclEi), a tool that automates the identification of toroidal nuclei, minimizing false positives while highly agreeing with the manual quantifications. Additionally, PIQUE identifies micronuclei for a convenient comparison of both CIN biomarkers. PIQUE is an open-source ImageJ plugin with a user-friendly interface that enables a wide scientific community to easily study the novel toroidal nucleus for the analysis of CIN levels in cellular models.

**Availability:** The plugin, the user manual, and the images used for the evaluation of PIQUE are available upon request from the authors.

## 1 Introduction

Cancer cells often exhibit large chromosomal alterations, as found in 80% of solid tumors (Hoevenaar *et al*., 2020). Chromosomal instability (CIN) is a hallmark of cancer, and understanding the mechanisms of chromosome missegregation is a key focus of cancer research. However, analysis of CIN in mitotic cells is challenging since mitosis is short and dividing cells make only 10% of the cell population. Therefore, biomarkers detectable in interphase cells are convenient for the study of CIN. Although chromosome missegregation can result in different aberrant nuclear morphologies (Gisselsson *et al*., 2001), the presence of micronuclei is usually the only phenotype considered to score for CIN level (Fig. 1A) (Krupina *et al*., 2021). Thus, complementary biomarkers are needed to obtain an exhaustive analysis of CIN status in cells. We recently identified the toroidal nucleus as a novel CIN biomarker (Almacellas *et al*., 2021), which is characterized by a doughnut-like shape with a void encompassing cytosolic components (Fig. 1A). Toroidal nuclei have been so far detected in 80% of the screened cell lines (Almacellas *et al*., 2021), and also in *S. cerevisiae* (Garcia *et al*., 2022), highlighting its potential value as readout across eukaryotes.

**Figure 1.**
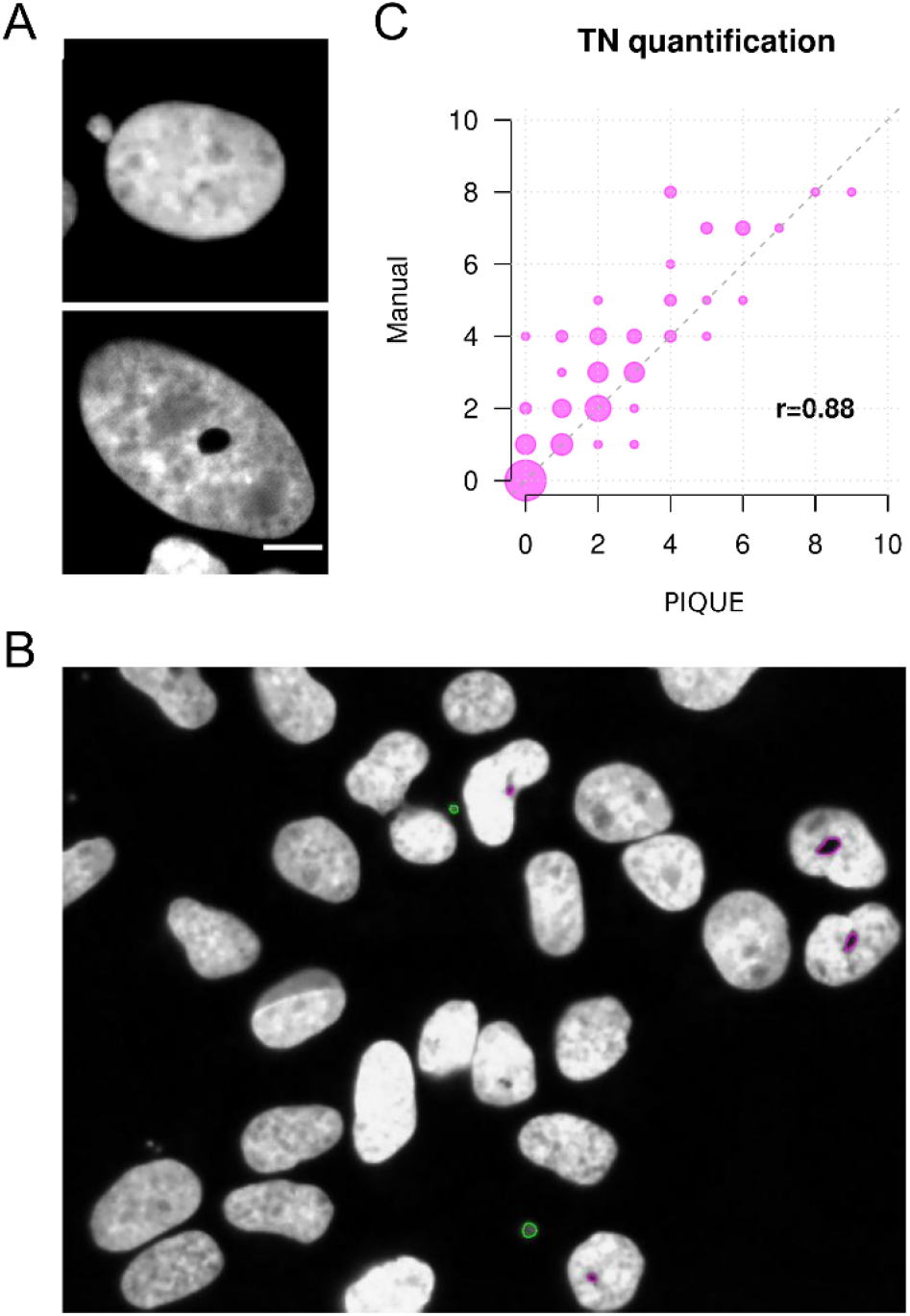
A) Cell nucleus with a micronucleus (top) and a toroidal nucleus (bottom). B) Representative output image generated by PIQUE in which four toroidal nuclei (in magenta) and two micronuclei (in green) were identified. C) Scatter plot showing the number of toroidal nuclei quantified by visual inspection and by PIQUE, and their Pearson’s correlation coefficient.

Even if most cancer cell lines exhibit an increase in toroidal nuclei and micronuclei (Almacellas *et al*., 2021; Krupina *et al*., 2021), the specific stimuli leading to the formation of these phenotypes are mostly unknown and might differ (Almacellas *et al*., 2021). Characterization of these stimuli requires extensive comparisons across cell types and conditions. Also, the prevalence of the toroidal nucleus is not fully understood, and the existing collections of morphological images, both in human, like the Broad Institute image collection (Ljosa *et al*., 2012), and yeast (Ohnuki *et al*., 2022; Mattiazzi Usaj *et al*., 2020), represent an invaluable resource to scan for its presence in very heterogeneous genetic backgrounds and conditions. Thus, further characterization of the toroidal nucleus demands the processing of large sets of images. However, quantification of toroidal nuclei currently relies on the manual analysis of fluorescent biological images (Pons *et al*., 2022), which is work-intensive and a limiting factor for large-scale analyses.

Here, we present PIQUE, an ImageJ plugin, open-source and freely available, for the efficient and reliable quantification of toroidal nuclei in large sets of fluorescent images.

## 2 Methods

U2OS and HeLa cells were grown on sterile coverslips and fixed with 4% PFA for 10 minutes before counterstaining with DAPI for DNA detection, as explained previously (Pons *et al*., 2022). We acquired 100 images with a fluorescent microscope equipped with a 405 nm laser and a 40x air objective, containing nuclei of heterogeneous size and shape, with different levels of intensity saturation depending on their cell cycle phase when fixed. We visually inspected the images and quantified the number of toroidal nuclei and micronuclei.

We automated the quantitative analysis of toroidal nuclei by implementing PIQUE, an ImageJ (Schneider *et al*., 2012) plugin to process biological images containing DAPI-stained nuclei. PIQUE identifies nuclei by applying a dynamic intensity threshold, which accounts for the specific saturation level in each biological image. To identify toroidal nuclei, which should manifest as a single void with an intensity similar to the background, PIQUE implements a single cell-based approach. After performing nuclei segmentation, it applies an intensity threshold to each nucleus proportional to the background and local intensities, disregarding nuclei with multiple low intensity regions. The cell-based analysis allows the detection of brighter toroidal nuclei in saturated cells, and penalizes unexposed DAPI-stained nuclei. For the identification of micronuclei, PIQUE locates small circular particles and calculates the distance to their closest cell nucleus centroid. Particles far from any nucleus, such as isolated debris, are thus discarded. Also, multiple micronuclei close to the same nucleus, which probably originate from the same CIN event, are considered only once. Size and circularity ranges for nucleus, toroidal nuclei, and micronuclei are user-defined parameters. It should be noted that PIQUE will be used for the screening of large collections of images in order to identify cell lines and conditions favoring the formation of toroidal nuclei. Thus, our approach and choice of parameters is inherently conservative, maximizing precision in order to minimize wrong calls that could compromise downstream analyses and conclusions.

PIQUE runs on single images and also on sets of images within a selected folder. For each run, PIQUE generates an output image highlighting all the identified relevant particles colored by their phenotype (Fig. 1B), including a header with the corresponding quantifications which are also reported in a separate text file. Importantly, PIQUE provides a spreadsheet per input image with topological descriptors of each identified nucleus, and whether they present any of the biomarkers, enabling future studies on morphological features of toroidal nuclei and micronuclei. If run on multiple images, PIQUE generates an additional summary spreadsheet with the corresponding quantifications.

## 3 Results

We ran PIQUE on a set of 100 biological images (Supplementary material), each consisting of 20 to 100 cells from different experiments in U2OS and HeLa cells. We manually quantified the images and found the number of toroidal nuclei and micronuclei to be highly correlated with PIQUE quantifications (Pearson’s correlation = 0.88 and 0.84, respectively; see Fig. 1C). We individually verified each of the 188 toroidal nuclei identified by PIQUE and found 166 to be correct (precision = 0.88) with a recall of 0.67. Precision and recall for micronuclei identification were 0.99 and 0.60, respectively.

The average running time per image was 0.5 seconds on a laptop with a i7-8550U processor. We successfully tested PIQUE with ImageJ version 1.53t in Ubuntu and Windows10, respectively, using the set of 100 biological images mentioned above.

## 4 Conclusions

PIQUE is a computational image processing tool which enables the unbiased and reproducible identification of toroidal nuclei, representing a substantial improvement compared to the current time-consuming manual quantifications prone to human bias and error. Being the first automated approach for the identification of toroidal nuclei, PIQUE offers a unique opportunity for further understanding the mechanisms involved in chromosome missegregation and CIN, both specifically relevant in cancer research. In all, PIQUE enables the optimal detection of nuclear biomarkers to assess CIN in eukaryotic cells and provides an effective tool for genotoxicity screens.

## References

Almacellas, E. et al. (2021) Lysosomal degradation ensures accurate chromosomal segregation to prevent chromosomal instability. Autophagy, 17, 796–813.

Garcia, M. et al. (2022) Nuclear ingression of cytoplasmic bodies accompanies a boost in autophagy. Life Sci Alliance, 5.

Gisselsson, D. et al. (2001) Abnormal nuclear shape in solid tumors reflects mitotic instability. Am. J. Pathol., 158, 199–206.

Hoevenaar, W.H.M. et al. (2020) Degree and site of chromosomal instability define its oncogenic potential. Nat. Commun., 11, 1501.

Krupina, K. et al. (2021) Causes and consequences of micronuclei. Curr. Opin. Cell Biol., 70, 91–99.

Ljosa, V. et al. (2012) Annotated high-throughput microscopy image sets for validation. Nat. Methods, 9, 637.

Mattiazzi Usaj, M. et al. (2020) Systematic genetics and single-cell imaging reveal *widespread morphological pleiotropy and cell-to-cell variability*. Mol. Syst. Biol., 16, e9243.

Ohnuki, S. et al. (2022) High-throughput platform for yeast morphological profiling predicts the targets of bioactive compounds. NPJ Syst Biol Appl, 8, 3.

Pons, C. et al. (2022) Detection of Nuclear Biomarkers for Chromosomal Instability. Methods Mol. Biol., 2445, 117–125.

Schneider, C.A. et al. (2012) NIH Image to ImageJ: 25 years of image analysis. Nat. Methods, 9, 671–675.

